# Leaf excision has minimal impact on photosynthetic parameters across crop functional types

**DOI:** 10.1101/2023.04.25.538279

**Authors:** John N. Ferguson, Tamanna Jithesh, Tracy Lawson, Johannes Kromdijk

**Affiliations:** Department of Plant Sciences, University of Cambridge, Cambridge, Cambridgeshire, CB2 3EA, UK; School of Life Sciences, University of Essex, Wivenhoe Park, Colchester, Essex CO4 3SQ, UK; Institute for Genomic Biology, University of Illinois at Urbana-Champaign, Urbana, Illinois, 61801, USA

**Author notes:** Current address: Centre for Crop and Environmental Science, Harper Adams University, Edgmond, Shropshire TF10 8NB, UK. Equal contribution (John N. Ferguson, Tamanna Jithesh).

**Keywords:** Leaf excision, photosynthesis, leaf reflectance, chlorophyll fluorescence, *Solanum lycopersicum*, *Hordeum vulgare*, *Zea mays*

## Abstract

Photosynthesis is increasingly becoming a recognised target for crop improvement. Phenotyping photosynthesis-related traits on field-grown material is a key bottleneck to progress here due to logistical barriers and short measurement days. Many studies attempt to overcome these challenges by phenotyping excised leaf material in the laboratory. To date there are no demonstrated examples of the representative nature of photosynthesis measurements performed on excised leaves relative to intact leaves in crops. Here, we tested whether standardised leaf excision on the day prior to phenotyping impacted a range of common photosynthesis-related traits across crop functional types using tomato (C3-dicot), barley (C3-monocot), and maize (C4-monocot). Potentially constraining aspects of leaf physiology that could be forecasted to impair photosynthesis in excised leaves, namely leaf water potential and abscisic acid accumulation, were not different between intact and excised leaves. We also observed non-significant differences in spectral reflectance and chlorophyll fluorescence traits between the treatments across the three species. However, we did observe some significant differences between gas exchange and photosynthetic capacity associated traits across all three species. This study represents a useful reference for those who perform measurements of this nature and the differences reported should be considered in associated experimental design and statistical analyses.

**Highlight:** Across the main photosynthesis functional types (C_3_-dicot, C_3_-monot, and C_4_-monot) of major crops (tomato, barley, and maize), measurements of photosynthetic parameters demonstrate few, but important, differences when measured on excised relative to intact leaves.

## Introduction

The global demand for food is expected to double by the middle of this century. Human population growth means that the rate of increase in food demand outpaces the annual rate of increase in crop productivity (Zhu *et al*., 2010; Kromdijk and Long, 2016; Lenaerts *et al*., 2019). This mismatch highlights the necessity to adopt novel crop improvement targets to achieve food security (Gojon *et al*., 2023). As the primary determinant of biomass accumulation, photosynthesis represents a sensible target to this end. Evidence from free air CO_2_ enrichment experiments has highlighted how increasing CO_2_ assimilation can lead to improved crop yields (Ainsworth and Long, 2021). Similarly, transgenic improvements to independent aspects of photosynthetic biochemistry have demonstrated that increasing carbon fixation is a realistic strategy for increasing yield (Driever *et al*., 2017; Yoon *et al*., 2020; De Souza *et al*., 2022).

Targeting photosynthesis in crop improvement efforts necessitates the ability to screen many hundreds or thousands of genotypes for photosynthesis-associated traits. This is important for quantifying variation to incorporate in selection models. Additionally, it is necessary for facilitating forward genetics to identify novel genes or genetic regions underlying said variation to target through molecular approaches, such as marker-assisted breeding or direct manipulation through genome editing. This is challenging because the gold-standard methods through which many photosynthesis-associated traits are phenotyped are often logistically challenging to perform in field trial-like environments and they are time consuming (Long, 2003; Walker *et al*., 2018). Standard methodologies, such as the commonly performed photosynthesis-CO_2_ response (A-*c_i_*) measurement used for modelling photosynthetic capacity can take up to 45-60 minutes to perform per sample. This is problematic because it limits the number of measurements that can be performed per day. This is especially true in the field relative to controlled conditions, where diurnal weather effects on physiological patterns, such as decreasing leaf water potential, photosystem II efficiency, and chloroplastic inorganic phosphate concentration, can transiently impact photosynthetic activity and the length of the measurement day (Murchie *et al*., 1999; Leakey *et al*., 2006). Logistical challenges arise, for example, due to specialised equipment not being suitable for use within crop canopies and because of the lack of access to power supplies.

A potential means through which to overcome some of these confounding effects and logistical challenges is to excise leaf material from plants of interest and phenotype those leaves in a more amenable and stable laboratory setting. Studies on woody species have shown that phenotyping photosynthetic- and reflectance-associated traits on the leaves of excised branches generates data that is highly comparable to that generated through phenotyping leaves on branches still attached to the tree (Richardson and Berlyn, 2002; Miyazawa *et al*., 2011; Akalusi *et al*., 2021). Over the past decade, a similar approach has routinely been employed for multiple crop species (e.g., Driever *et al*., 2014; Ferguson *et al*., 2021; Montes *et al*., 2022). Here, leaves, or tillers harbouring a leaf of interest, are excised from plants and the cut end is placed into water in an attempt to maintain hydraulic conductivity. For photo-physiological phenotyping, the maintenance of hydraulic efficiency is important for replacing water lost via transpiration (Müllers *et al*., 2022), which is instrumental to keep stomata open for CO_2_ uptake. It is possible that the demonstrable success of employing this approach with woody species is at least in part due to the presence of secondary xylem. Secondary xylem facilitates water transport, but it is also critical in protecting the primary xylem from mechanical damage (Miodek *et al*., 2021), such as that that might ensue after branch excision. This presence of secondary xylem is a key anatomical difference that distinguishes the tree species where this approach has been demonstrated to work as effectively as *in-situ* phenotyping and the herbaceous annual crop species for which this technique is now regularly employed. Due to this key difference and others, it is plausible that the mechanistic basis that underpins the success of performing photosynthesis measurements on foliage of detached branches of woody species does not necessarily translate to herbaceous species. Consequently, a key knowledge gap exists here with respect to understanding the extent to which measurements made on excised leaves are representative of those made on intact leaves for crop species.

Addressing the above-described knowledge gap is a high-priority due to the rapid uptake of this approach for phenotyping photosynthesis in crop species. Variations on this technique have been reported in studies involving many key crop species, however we are not aware of any studies that employ this approach and show that the data they obtain are representative of what would have been obtained from intact material. Information to this end will be key for emphasising the value of this approach. Maize is perhaps the species with which this approach has been most employed. For example, Markelz *et al*., (2011) excised leaves from maize plants growing in a free air CO_2_ enrichment (FACE) experiment and under variable soil nitrogen availabilities to perform *A*-*c_i_* response measurements to test interactive effects of CO_2_ and nitrogen supply on photosynthetic capacity. Similarly, Liao *et al*., (2022) phenotyped excised maize leaves from plants growing under full- and deficit-irrigation to again perform *A*-*c_i_* response measurements to test how water availability influenced photosynthetic capacity and stomatal limitations to photosynthesis. As well as phenotyping small numbers of maize genotypes to test environmental interactions (See also: Leakey *et al*., 2006) and Choquette *et al*., (2019)), this approach has also been employed for maize to facilitate screening of large populations for forward genetics purposes. For example, Xie *et al*., (2021) used this approach to phenotype a mapping population consisting of 197 recombinant inbred lines (RILs) for light-saturated gas exchange. By utilising this approach, this study was able to identify genetic regions underpinning variation in photosynthesis and associated traits. An identical approach has also been utilised for phenotyping genetic diversity in sorghum. Here, this approach enabled by-far the largest survey of genetic variation in photosynthesis, where over 800 genetically distinct sorghum varieties were characterised enabling genome- and transcriptome-wide association mapping (GWAS and TWAS) to pinpoint precise genes underlying variation in photosynthesis (Ferguson *et al*., 2021). Phenotyping photosynthesis of excised leaves has also been used in other C_4_ species, such as Miscanthus, where this approach has facilitated testing of the adaptation of photosynthesis to self-shading (Collison *et al*., 2020), and sugarcane, where it has facilitated testing of the response of photosynthesis to the application of sugars (McCormick *et al*., 2008).

Despite C_4_ crops typically having more physiologically robust leaves, measuring photosynthesis in excised material has also been performed on studies focusing on monocot and dicot C_3_ crops. For example, excised tillers were used to phenotype flag-leaves for a suite of photosynthesis-related traits in a diversity set of 64 wheat accessions at different developmental stages (Driever *et al*., 2014; Carmo-Silva *et al*., 2017). This approach has also been used in barley where the leaf of interest was directly excised at its base rather than harvesting the entire tiller to test how direct nitrate feeding impacted photosynthesis and sugar synthesis (Morcuende *et al*., 2005). In rice, excised leaf material has also been used to measure chlorophyll fluorescence across distinct varieties to facilitate GWAS of fluorescence parameters (Ferguson *et al*., 2020). Similar fluorescence experiments on excised leaf segments of wheat provided some preliminary evidence for an impact of the time between excision and measurement, where a minimal waiting time may be needed to properly relate observations to differences in photosynthetic efficiency (McAusland *et al*., 2019). To the best of our knowledge, utilisation of excised leaf material in C_3_ dicots is constrained to studies on soybean. For example, Morgan *et al*., (2004) cut petioles of fully expanded leaves of soybean plants growing in an open-air elevated ozone experiment to perform chlorophyll fluorescence, A-*c_i_* and photosynthesis light-response (AQ) measurements to understand the photosynthetic response to ozone stress. The same technique was operated by Montes *et al*., (2022) to perform *A*-*c_i_* measurements and concurrent measurements of hyperspectral reflectance to build models for predicting photosynthetic capacity in the field and for facilitating GWAS.

Excision plausibly elicits an abscisic acid (ABA) biosynthesis response in leaves in a manner similar to that which promotes the stabilisation of leaf water status (Beardsell and Cohen, 1975) and/or resistance to herbivory (Vos *et al*., 2013, 2019; Nguyen *et al*., 2016). This is an important consideration here since ABA accumulation within leaves also serves to close stomatal apertures and reduce transpirational water loss in response to drought, which consequently impairs carbon uptake (Blatt, 2000; Schroeder *et al*., 2001). Further to this, there may well be a myriad of minor-to-major alterations of leaf properties, e.g., changes to xylem tension, cell turgor, leaf water potential, pigmentation, etc. (Govindjee *et al*., 1981; Carter, 1991; Canny, 1997; Richardson and Berlyn, 2002; Tyree and Cochard, 2003), that could impact the photosynthetic activity of excised leaves. Due to the reality of these potential pitfalls and because of our reliance on performing photosynthesis measurements on excised leaf material in crop research, it is critical to understand whether leaf excision impacts photosynthesis in crop species. To this end and with this study we tested how standardised leaf excision on the day before phenotyping impacted light-saturated photosynthesis, photosynthetic capacity, chlorophyll fluorescence, hyperspectral reflectance, ABA accumulation and leaf water potential. For this we employed reference genotypes of tomato, barley, and maize, which represent the main photosynthesis functional types of crops, i.e., C_3_ monocot, C_3_ dicot and C_4_ monocot.

## Materials and methods

### Plant Material and Experimental Design

Reference genotypes for tomato, barley and maize were used for this study. These genotypes were M83 (tomato), Golden Promise (barley), and B73 (maize). All plants were grown at the Plant Growth Facility of the University of Cambridge. Seeds from all species were sown into modular trays before being transplanted into larger pots once the seedlings were established. Barley and tomato were grown in 1.5L sized pots. Maize was grown in 6L sized pots with two plants per plot. Barley was grown in multipurpose compost (M3 Compost, Levington Ltd., UK). Tomato was grown in a seedling and cuttings compost (F2 Compost, Levington Ltd., UK) and fertilised with specialised tomato feed (Tomorite, Levington Ltd., UK). Maize was grown in a 2:1 mix of these two compost types with added perlite. Tomato and maize were grown in the same growth room where the photoperiod was set to 14 hours. The light level was set to 600 µmol m^-2^ s^-1^ photosynthetically active radiation (PAR). The humidity was set to 65% relative humidity (RH). The air temperature was set to 28°C in the daytime and 20°C in the night-time. The growth room of the barley was set to a 16-hour photoperiod. The light level was set to 400 µmol m^-2^ s^-1^ PAR. The humidity was set to 60% RH. The air temperature was set to 22°C in the daytime and night-time.

Prior to any given measurement day, plants were transferred from the Plant Growth Facility to the laboratory at the Department of Plant Sciences (University of Cambridge) at 14.00 h. All plants were checked to be well watered and left on the laboratory bench. At 17.00 h, those plants that were to be measured as excised were cut. For tomato, the youngest fully expanded leaf was cut at the petiole below the two leaflets adjacent to the terminal leaflet. The petiole was then immediately placed under water and recut before being transferred to a 15 ml falcon tube under water. For barley, the youngest fully expanded leaf was excised at its base. The leaf was then recut just above the initial excision under water and transferred under water to a 15ml falcon tube. For maize, the youngest fully expanded leaf was excised at its base, recut just above the initial excision under water and transferred under water to a 50ml falcon tube. We ensured that all falcon tubes contained sufficient water so that the excision point was consistently submerged. Excised leaves were left on the same lab bench as their intact counterparts.

On the measurement day, excised and intact leaves were subject to a series of parallel measurements as highlighted in Supplemental Figure S1, with details provided below. Tomato plants were measured eight weeks post sowing across a two-day period. Barley plants were measured across two separate two-day periods one week apart, four and five weeks post sowing. Maize plants were measured across two separate two-day periods two weeks apart, six and eight weeks post sowing.

### Leaf-level gas exchange

Leaf-level gas exchange measurements were performed using two LI-COR 6400XT infra-red gas analysers fitted with 6400-40 fluorometer LED light sources (LI-COR Biosciences, Lincoln, NE). To measure light-saturated gas exchange, conditions within the gas exchange chamber were set as follow: 25°C block temperature; 400 µmol s^-1^ air flow; 65-75% relative humidity (RH); 400 µmol s^-1^ reference CO_2_ concentration; 1500 µmol m^-2^ s^-1^ PAR. Once *A_N_* and *g_S_* were stable, light saturated rates of gas exchange were logged before initiating a CO_2_ response curve as described below.

For the tomato and barley CO_2_ response curves, all conditions were kept as described above with gas exchange being measured at a series of reference CO_2_ concentrations as follows: 400, 300, 200, 100, 50, 400, 400, 700, 1000, 1300, and 1800. Rates of gas exchange were logged at each point according to stability criteria waiting a minimum of 90 seconds and a maximum of 120 seconds at each point.

For the maize CO_2_ response curves, all conditions were again kept as described for the light saturated gas exchange. Reference CO_2_ was then increased in the following steps: 400, 600, 800, 1000, and 1250. Reference CO_2_ was then returned to 400 and rates of *A_N_* and *g_s_* were allowed to re-stabilise before reducing reference CO_2_ in the following sequence: 400, 300, 250, 200, 100, 75, and 25. Rates of gas exchange were logged at each point according to stability criteria waiting a minimum of 60 seconds and a maximum of 90 seconds at each point.

For tomato and barley, *A_N_*-*c_i_* data from the response curves were fit according to the FvCB model (Farquhar *et al*., 1980) using the fitacis() function from the R package plantecophys (Duursma, 2015). This was achieved using the bilinear method to estimate the transition point. From this, estimates of the maximum rate of Rubisco carboxylation (*V_cmax_*) and the maximum rate of electron transport for RuBP regeneration (*J_max_*) on a *C_i_* basis were obtained. For maize, *A_N_*-*c_i_* data from the response curves were fit using a custom R function following von Caemmerer, (2000). Here, the initial relationship between *A_N_* and *c_i_* was used to estimate the maximum rate of carboxylation by PEPC (*V_pmax_*). Additionally, the horizontal asymptote of a four-parameter non-rectangular hyperbola was used as an estimate of CO_2_- and light-saturated photosynthesis (*V_max_*).

### Hyperspectral reflectance

Hyperspectral reflectance measurements were performed using an ASD FieldSpec 4 Standard-Res Spectroradiometer equipped with a leaf-clip (Malvern Panalytical, Malvern, UK). The light source of the FieldSpec was allowed to warm up for 45 minutes prior to measurements. Hyperspectral reflectance was always measured on the same area of the leaf that gas exchange was performed on and always with the light source pointing towards the adaxial leaf surface. For each leaf, three technical replicates of hyperspectral reflectance were performed within the spot used for gas exchange measurements. Reflectance at each wavelength was then averaged across the three technical replicates for further data analyses. These measurements were performed immediately following the CO_2_-response gas exchange measurements.

### Chlorophyll fluorescence

Chlorophyll fluorescence measurements were performed using both the previously described LICOR 6400XT leaf chamber fluorometer and using a closed chlorophyll fluorescence imaging system (FluorCam FC 800-C, PSI, Czech Republic). For both, a similar program was designed to measure the induction of non-photochemical quenching (NPQ) in response to actinic light and the subsequent relaxation of NPQ following the actinic light being switched off.

Following the afternoon hyperspectral reflectance measurements, a portion of leaf was dark adapted using aluminium foil for 45 minutes. After dark adaption, leaves were clamped within the gas exchange chamber where conditions were as described previously except the light source was switched off. Following estimation of dark-adapted maximum fluorescence (*F_m_*), the light source was switched on to 1500 µmol m^-2^ s^-1^ PAR for 10 minutes and a series of 12 saturating pulses were performed to estimate maximal fluorescence (*F_m_*’) at different time- points. The light source was then subsequently switched off for 12 minutes and a further series of eight saturating pulses were performed again to estimate *F_m_*’. NPQ at each point of measurement of chlorophyll fluorescence was then calculated as (*F_m_*-*F_m_*’)/*F_m_*’.

Samples for measurements of chlorophyll fluorescence in the closed system were first analysed for leaf water potential using a Scholander pressure chamber (see below). For barley and maize, leaf strips approximately 3cm x 5cm were taken from the middle portion of the measured leaf and for tomato the same sized strip was from the measured terminal leaflet. Leaf strips were arranged onto damp filter paper and encased between non-reflective glass as per Ferguson *et al*., (2020). These samples were then dark adapted overnight and measured for NPQ induction and relaxation within the closed chlorophyll fluorescence system following the same sequence of steps as described when using the LI-COR 6400 leaf chamber fluorometer.

Using the NPQ data generated by both the leaf chamber fluorometer and the closed fluorescence system, separate exponential models were fit to the light induction (Equation 1) and dark relaxation (Equation 2) components.

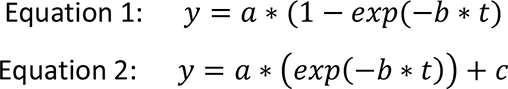

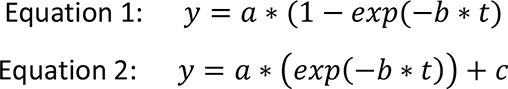

For both equations *a* represents the amplitude of the induction or relaxation response (a_ind and a_rel), *b* is the rate constant for the induction or relaxation of NPQ (b_ind and b_rel), *c* is an offset to account for a non-zero intercept (relaxation only), *t* is the measurement time point, and *y* is NPQ.

### Leaf water potential

Measurements of leaf water potential were performed using a Scholander pressure chamber (Model 600, PMS Instrument Company, OR, USA) following manufacturer instructions. For intact leaves, a clean excision was made with a razor blade towards the base of the leaf for maize and barley or the petiole for tomato before immediately sealing the leaf into the chamber to measure the leaf water potential. For excised leaves, a new additional excision was made 3cm above the cut from the previous day before immediately measuring leaf water potential in the same manner.

### ABA analysis

Separate plants were grown for analyses of leaf ABA concentration. Here, the plants were grown, and leaves were excised in exactly the same manner as described in the “Plant Material and Experimental Design” section. Barley and maize plants were measured five weeks post sowing and tomato plants were measured eight weeks post sowing. Samples for ABA concentration measurements were taken in the morning (10am). Each sample was taken from the middle portion of the leaf or terminal leaflet taking care to avoid the midrib in barley and maize. Samples were immediately flash frozen and subsequently analysed for ABA concentration using a competitive ELISA kit (Phytodetek Immunoassay Kit for ABA, Agdia, Elkhart, IN) following the manufacturer’s instructions. The area of each sample was measured before flash freezing, so that ABA concentration could be expressed on leaf area basis.

### Data processing and statistical analyses

All statistical analyses were performed within the R environment (Reference). All figures were generated using ggplot2 (Valero-Mora, 2010) and gridExtra with some pre-processing in Affinity designer (Serif, Nottingham, UK).

We performed two-way repeated measures analysis of variance (ANOVA) comparison of means tests to test for significance of the treatment (excised or intact), the time of measurement (AM or PM), and their interaction (treatment by time) for all gas exchange- associated traits. We performed one-sample t-tests to test for differences in leaf water potential and the chlorophyll fluorescence associated traits between excised and intact leaves for each species independently. The chlorophyll fluorescence traits measured using the gas exchange leaf chamber fluorometer and the closed chlorophyll fluorescence system were analysed independently as instrumental differences in sample conditioning, excitation wavelength as well as detection fore optics preclude direct comparisons.

The asdreader R package was used for managing the hyperspectral reflectance data following best practise outlined in Burnett *et al*., (2021). Hyperspectral reflectance data were initially analysed through a spectra-wide approach where one-way ANOVA comparison of means tests were performed to test for differences in reflectance between excised and intact leaves at each wavelength from 250-3500nm. Separate tests were performed for each species and each time point, i.e., AM and PM. We calculated a Bonferroni significance threshold (α = 0.05) to avoid identification of false positive results. It is worth noting that since there is covariation between wavelengths of a given spectra, this threshold is likely to be somewhat stringent. For visualisation purposes, we transformed the p-value associated with each wavelength by - log_10_.

We additionally used the same hyperspectral reflectance data to calculate multiple reflectance indices (Supplemental Table S1). We used these indices to perform repeated measures two-way ANOVA comparison of means tests separately for each species and as described previously to test for effects due to treatment, timepoint, and their interaction.

## Results

### Leaf water potential and ABA accumulation

At the end of the measurement day, we measured leaf water potential of all measured leaves (Figure 1, Supplemental Figure S1). For tomato and maize, we did not observe a significant effect of leaf excision on leaf water potential. However, for barley the leaf water potential of excised leaves showed a small but significant difference with that of the intact leaves (-0.193 and -0.237 MPa, respectively; Figure 1b).

**Figure 1.**
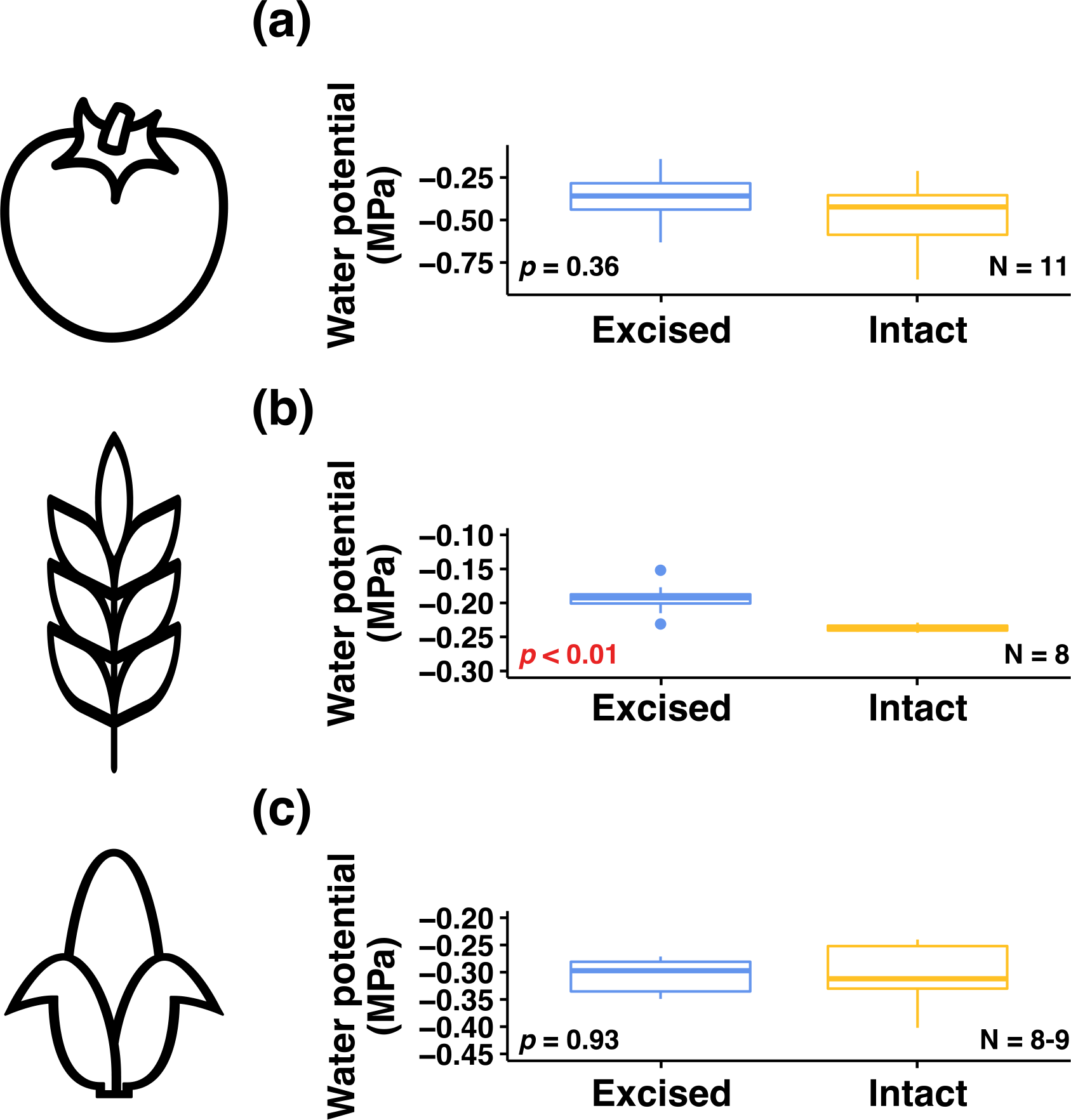
Boxplots showing differences in leaf water potential between intact and excised leaves for (a) tomato, (b) barley, and (c) maize. The p-values associated with the corresponding one-sample t-test are inset in each panel.

Leaf ABA concentration in excised and intact leaves was measured in the morning after the excision. For all three species, leaf excision did not result in a significant up- or down- regulation in ABA accumulation relative to intact leaves at this timepoint (Figure 2).

**Figure 2.**
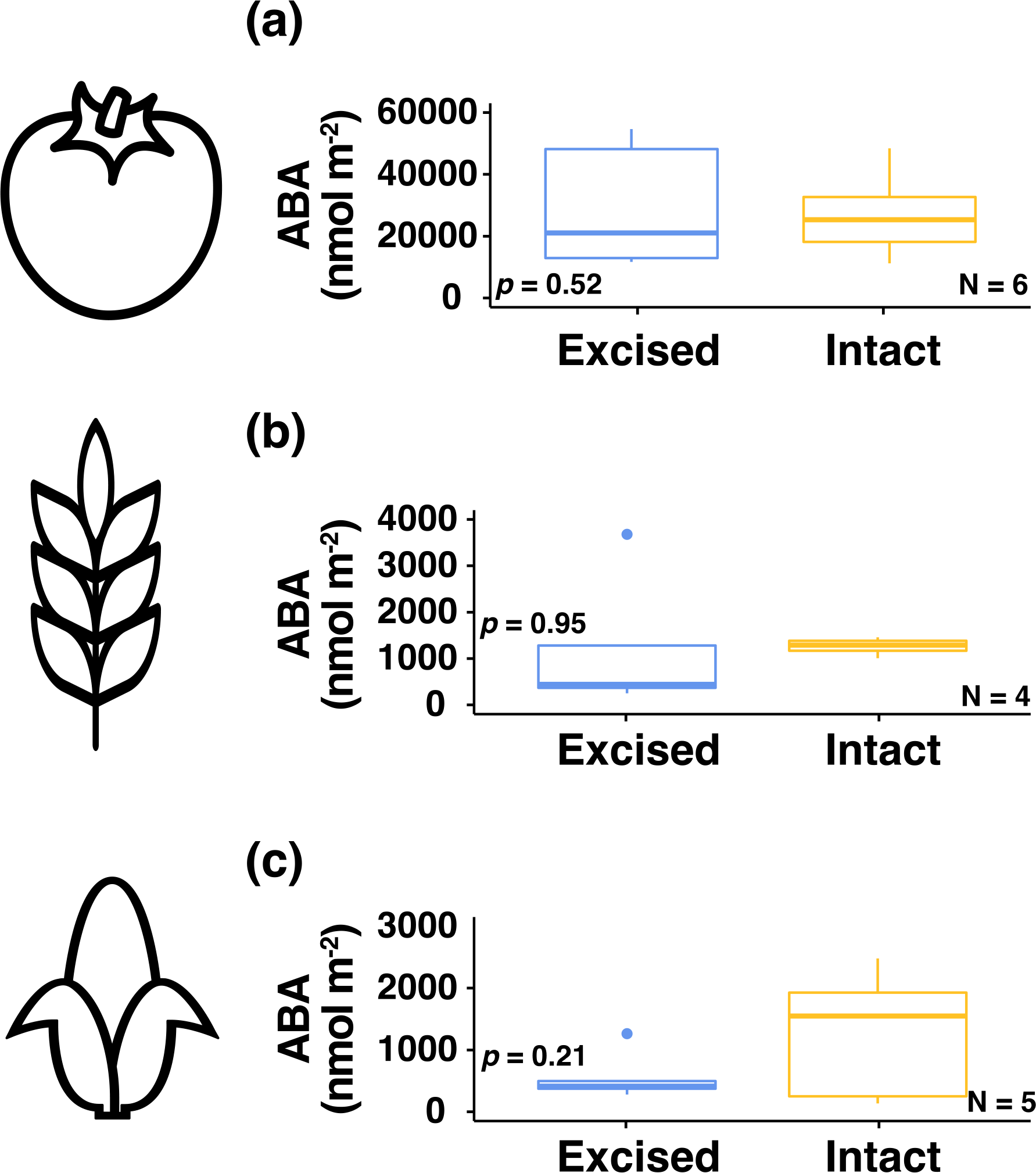
Boxplots showing differences in leaf ABA concentration between intact and excised leaves for (a) tomato, (b) barley, and (c) maize. The p-values associated with the corresponding one-sample t-test are inset in each panel.

### Light saturated gas exchange

Light-saturated gas exchange was logged prior to the initiation of the *A_N_*-*c_i_* response measurements in the morning and the afternoon (Supplemental Figure S1, Figure 3). Across all three species, leaf excision did not have a significant effect on *A_N_*. The timing of the measurement did have a significant effect on *A_N_* for barley and maize, but not tomato. Here, *A_N_* reduced in the afternoon for barley and maize. This reduction in *A_N_* was independent of the treatment, where *A_N_* declined in both intact and excised leaves (Figure 3e, i). Neither treatment, time, nor their interaction had a significant effect on *g_s_* for tomato (Figure 3b). Time had a significant effect on *g_s_* for barley, where it declined into the afternoon as per *A_N_* (Figure 3f). The excision treatment did have a significant effect on *gs* for maize, where it was reduced in excised leaves. Although the treatment by time interactive effect for *g_s_* was not significant, the difference between excised and intact leaves did appear to increase into the afternoon (Figure 3j). We did not detect a significant effect of treatment, time, or their interaction on *c_i_* for barley (Figure 3g). However, for tomato, *c_i_* increased significantly in the afternoon for excised and intact leaves (Figure 3c). For maize, all tested independent variables had a significant effect on *c_i_*. Intact leaves demonstrated higher *c_i_* than excised leaves in the morning and the afternoon, but the difference increased in the afternoon (Figure 3k). Significant effects on *iWUE* mirrored those observed for *c_i_*, such that it significantly declined in the afternoon for both tomato and maize, and it was significantly increased in excised leaves of maize (Figure 5d, l). With respect to the diurnal effect on *iWUE* and *c_i_* for maize, it is worth noting that the changes into the afternoon are much stronger in intact leaves, i.e., *iWUE* and *c_i_* are more stable across the day in excised leaves.

**Figure 3.**
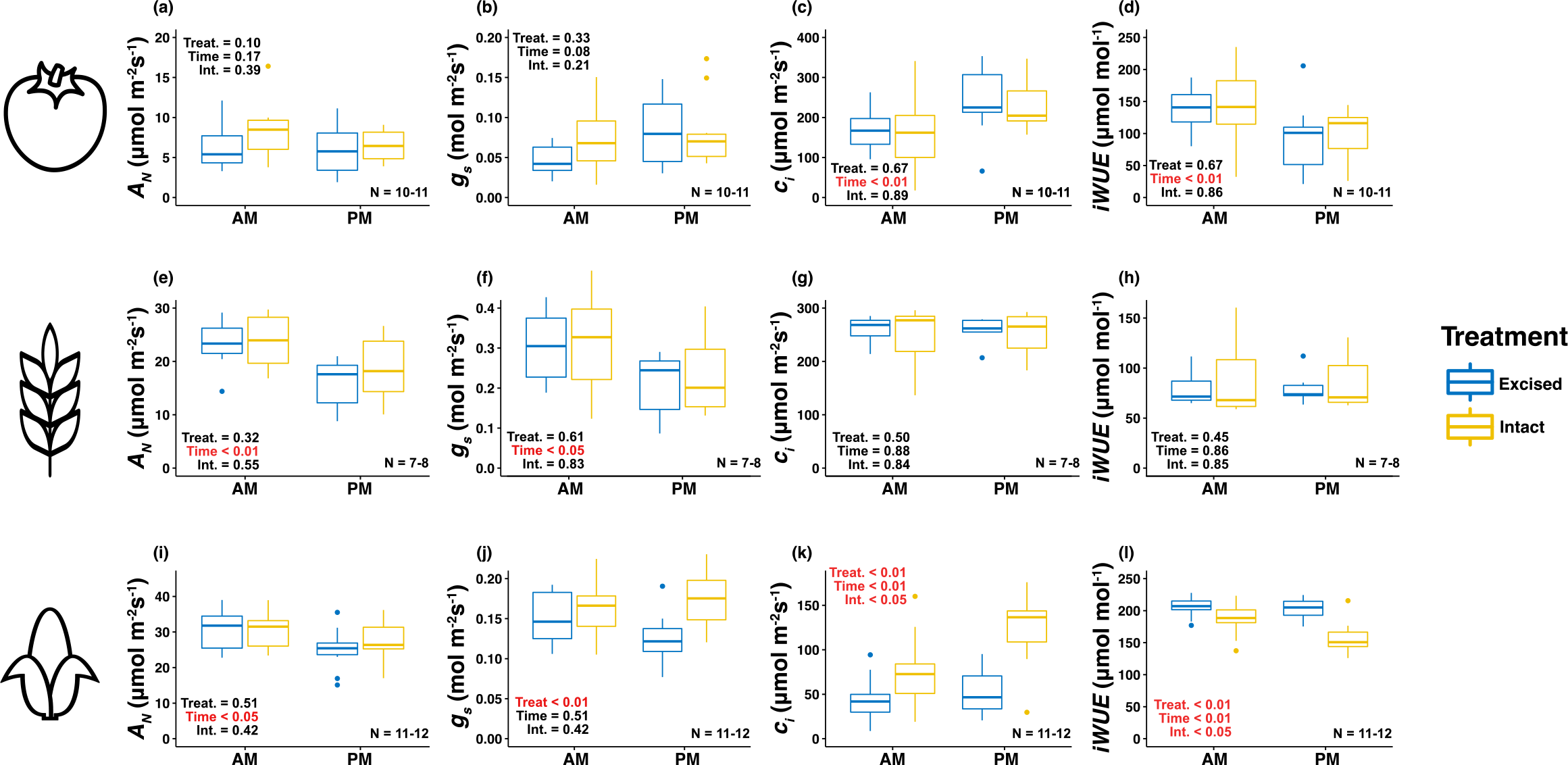
Boxplots showing differences in light-saturated gas exchange between excised and intact leaves in the morning (AM) and afternoon (PM) for tomato (a-d), barley (e-h), and maize (i-l). Net photosynthesis (*A_N_*), stomatal conductance (*g_s_*), the concentration of intracellular CO_2_ (*c_i_*), and intrinsic water use efficiency (*iWUE*) are shown. The p-values associated with the corresponding two-way analysis of variance (ANOVA) are inset in each panel.

### Photosynthetic capacity

We modelled *A_N_*-*c_i_* response measurements to estimate photosynthetic capacity in the morning and the afternoon of the three species (Supplemental Figure S1, Figure 4). Across the three species, time had a significant effect on all modelled parameters except *V_pmax_* in maize (Figure 4k). In all other cases, the photosynthetic capacity parameters declined into the afternoon regardless of whether the leaf being measured was excised or intact (Figure 4c, d, g, h, l). We only detected one significant treatment effect, where *V_cmax_* was significantly reduced in excised barley leaves, however the reduction appeared relatively minor (Figure 4g). In addition, the p-value associated with the treatment effect for *V_pmax_* in maize was close to the traditional 5% significance threshold (Figure 4k). For all traits and species, we did not observe a significant interaction effect of treatment and time, suggesting that any observed effects were temporally consistent (Figure 4).

**Figure 4.**
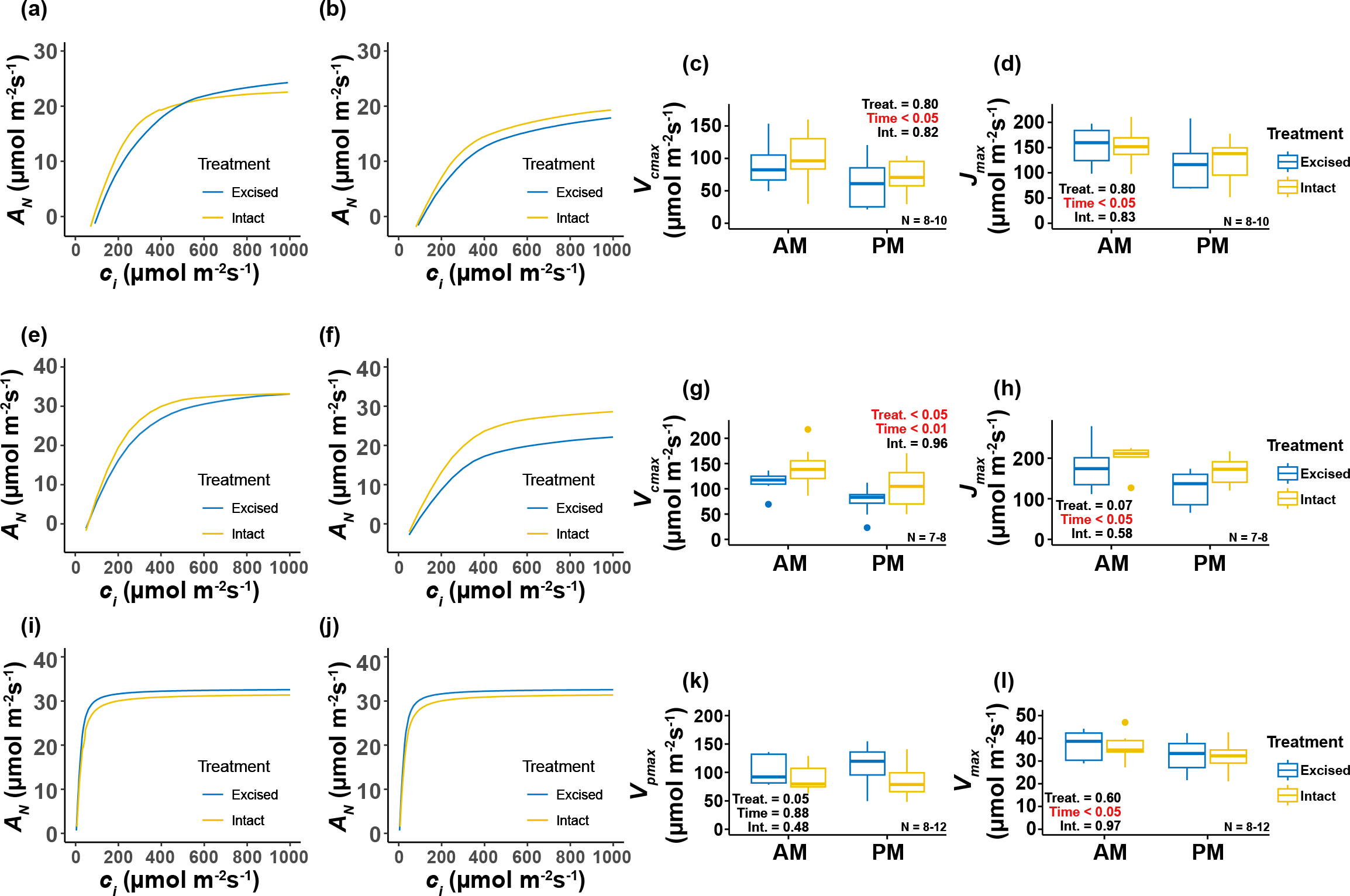
Differences in photosynthetic capacities modelled via photosynthesis-CO_2_ response measurements between intact and excised leaves. (a-b) The response of photosynthesis (*A_N_*) to increasing intracellular CO2 concentration (*c_i_*) in the (a) AM and (b) PM for tomato. The solid line represents the mean fit of the FvCB model and the shaded area represents the standard error of the mean. (c) The maximum rate of Rubisco carboxylation (*V_cmax_*) for tomato in the AM and PM. (d) The maximum rate of electron transport for RuBP regeneration (*J_max_*) for tomato in the AM and PM. (e-f) The response of *A_N_* to increasing *c_i_* in the (e) AM and (f) PM for barley. The solid line represents the mean fit of the FvCB model and the shaded area represents the standard error of the mean. (g) *V_cmax_* for barley in the AM and PM. (h) *J_max_* for barley in the AM and PM. (i-j) The response of *A_N_* to increasing *c_i_* in the (i) AM and (j) PM for maize. The solid line represents the mean fit of a four-parameter non-rectangular hyperbola and the shaded area represents the standard error of the mean. (k) The maximum rate of carboxylation by PEPc (*V_pmax_*) for maize in the AM and PM. (l) The maximum rate of photosynthesis (*V_max_*) for maize in the AM and PM.

### Leaf reflectance

Hyperspectral reflectance was measured following the *A_N_*-*c_i_* response measurements in the morning and the afternoon (Supplemental Figures S1, S2). None of the specific wavelengths showed any difference in reflectance between excised and intact leaves in the morning or the afternoon for any species according to the Bonferroni significance threshold (Figure 5). Despite the lack of significant wavelength-treatment associations, it is still interesting to observe the distinct species-specific patterns in spectra-wide reflectance differences between the treatments which can be interpreted based on the dominant factors that are known to control leaf reflectance across the hyperspectral range. The pattern of p-values across the spectral range were similar in the morning and the afternoon for tomato (Figure 5a-b) and maize (Figure 5e-f), however the pattern was distinctly different across these time points for barley (Figure 5c-d). For tomato, p-values showed a general trend of decreasing (inverse for - log_10_(p-values)) from 300 to 1800nm before increasing between 1800 and 1900nm. After this, they decreased to roughly what they were before the increase before increasing again toward the end of the spectral range (Figure 5a-b). This pattern suggests potential differences in leaf water status between the excised and intact tomato leaves. For maize, the pattern of p-values across the spectral range partially-resembled the pattern of reflectance (Figure 5e-f; Supplemental Figure S2). Here, there was a broad peak of low p-values between 600 and 1400nm and a narrower peak between 1500 and 1800nm. Here, the initial peak may reflect differences in leaf structure and the secondary peak is often associated with differences in leaf water status. For barley in the morning, the pattern of p-values across the spectral range was similar to that of maize reflecting potential effects of excision on leaf structure and water absorption (Figure 5c, e, f). However, the initial portion of the range was also characterised by comparatively reduced p-values, especially around 400nm (Figure 5c), which may suggest an effect on the absorption of light by chlorophyll b. In the afternoon, the broad peak of low p-values between 600 and 1400nm disappeared, but there were still minor peaks between 600 and 700nm and between 1800 and 2000nm (Figure 5d), suggesting potential minor effects on chlorophyll absorption and water status respectively.

**Figure 5.**
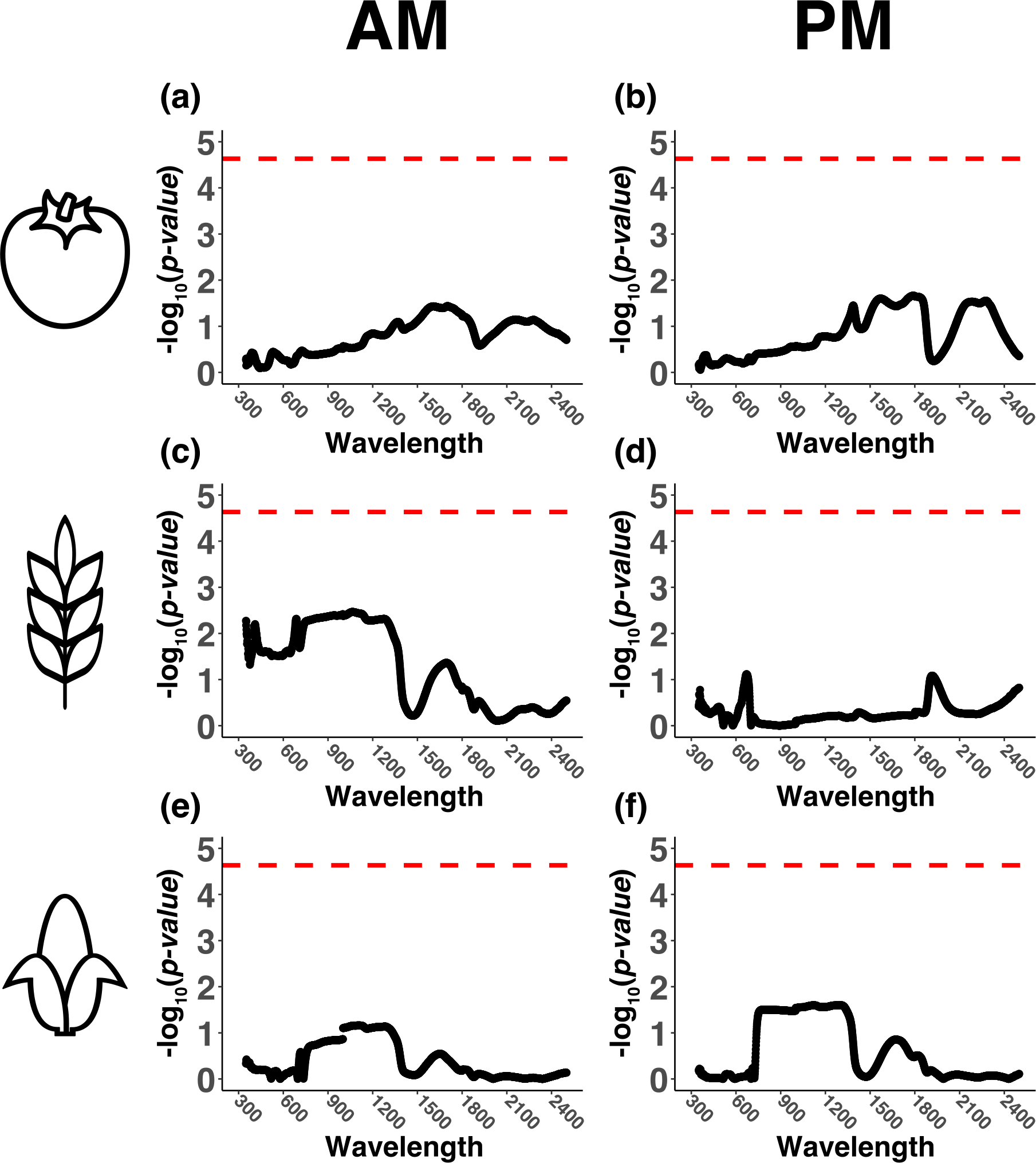
The effect of leaf excision on hyperspectral reflectance in the morning (AM) and afternoon (PM) for tomato, barley, and maize. (a) AM tomato. (b) PM tomato. (c) AM barley. (d) PM barley. (e) AM maize. (f) PM maize. Each panel shows the -log_10_ transformed p-value from a one-way analysis of variance (ANOVA) comparing reflectance of intact and excised leaves at each incremental wavelength from 350-2500nm. The dashed red lines indicate a Bonferroni significance threshold (α = 0.05).

Using the hyperspectral reflectance data (Supplemental Figure S2), we additionally calculated reflectance indices related to nitrogen status, chlorophyll content and leaf water status (Supplemental Table S1). We examined how these were affected by leaf excision and time of measurement, i.e., AM and PM (Supplemental Figure S1), via two-way repeated measures ANOVA tests as per the previously described photosynthesis traits (Supplemental Figures S4- 6). Although this approach did find some significant differences, the observed effects were minor in all cases.

For tomato, we did not observe any significant treatment, time, or treatment by time interactions for any of the reflection indices. (Supplemental Figures S3). For barley, we observed a significant time effect for the Datt Index, which is a proxy of total chlorophyll content, where it was higher in the afternoon (Supplemental Figure S4b). For barley, we also observed a significant interactive effect for the moisture stress index (MSI), which is an inverted proxy for water content, where excised leaves had a higher MSI in the morning only, which suggest their water content was reduced compared to intact leaves at this point but not again in the afternoon (Supplemental Figure S4d). For maize, the only significant effect observed was a treatment effect for the Datt Index, which was marginally higher in intact leaves compared to excised leaves suggesting chlorophyll content was reduced following leaf excision (Figure S5b) .

### Chlorophyll fluorescence

Chlorophyll fluorescence was measured using the leaf chamber fluorometer that was paired to the gas exchange platform and on the subsequent day using the closed chlorophyll fluorescence system (Supplemental Figure S1). We tested for significant differences between samples taken from excised and intact leaves for *F_v_*/*F_m_*, the rates of induction and relaxation of NPQ (b_ind and b_rel), and maximum NPQ. For the samples measured using the closed chlorophyl fluorescence system, we did not observe a significant difference for any of these four parameters (Figure 6). This was true for all of these same parameters as measured in the leaf chamber fluorometer, expect for tomato b_ind, which was observed to be significantly faster in intact versus excised leaves (Supplemental Figure S6b). It is worth noting that the p-values associated with the one-sample t-tests comparing excised and intact tomato leaves for *F_v_*/*F_m_* and maximum NPQ were close to the traditional 0.05 significance threshold (0.07 and 0.09 respectively), where intact leaves appeared to have higher photosystem II maximum efficiency and light adapted NPQ (Supplemental Figure S5b).

**Figure 6.**
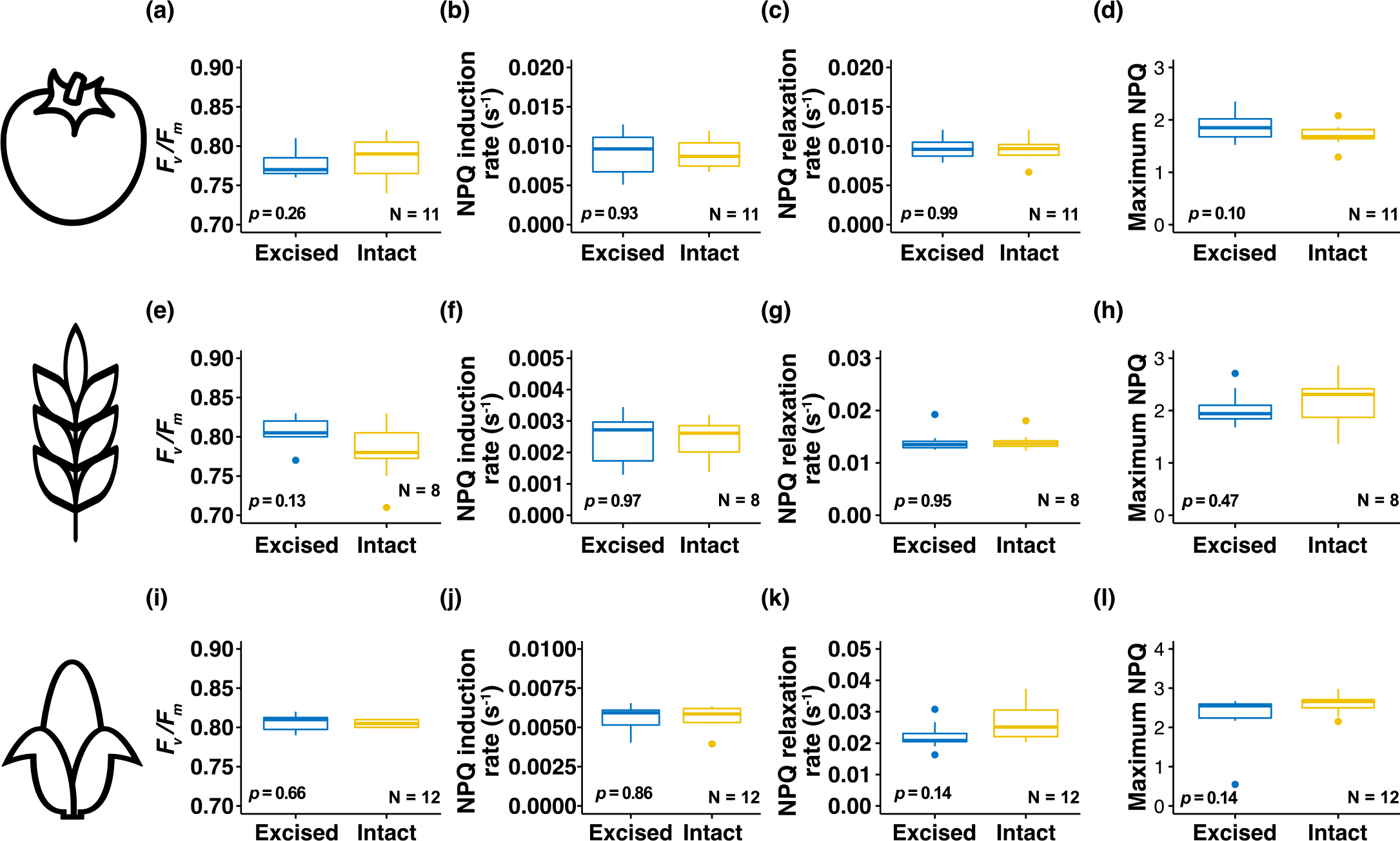
Boxplots showing variation in chlorophyll fluorescence-associated traits between excised and intact leaves measured in a closed chlorophyll fluorescence system. (a) Maximum efficiency of photosystem II (*F_v_*/*F_m_*) in tomato. (b) The rate of induction of non-photochemical quenching (NPQ) in tomato. (c) The rate of relaxation of NPQ in tomato. (d) Maximum NPQ in tomato. (e) *F_v_*/*F_m_* in barley. (f) The rate of induction of NPQ in barley. (g) The rate of relaxation of NPQ in barley. (h) Maximum NPQ in barley. (i) *F_v_*/*F_m_* in maize. (j) The rate of induction of NPQ in maize. (k) The rate of relaxation of NPQ in maize. (l) Maximum NPQ in maize. The p-values associated with the corresponding one-sample t-test are inset in each panel.

## Discussion

Screening variation in photosynthetic traits across large numbers of genotypes is a key requirement for targeting photosynthesis for crop improvement. Large-scale screening to this end is fraught with logistical challenges that are commonly circumvented via phenotyping excised leaf material (See: Introduction). A common limitation of these studies is the lack of evidence highlighting the representative nature of such phenotyping, which accordingly represents a key knowledge gap for this field. To this end, the principal aim of this study was to ascertain if photosynthetic parameters measured on excised leaves were representative of the same parameters measured on intact leaves in tomato, barley, and maize. There are at least three studies that have attempted to quantify potential differences in non-crop species. Using the Japanese stone oak (*Lithocarpus edulis*) evergreen tree species, Miyazawa *et al*., (2011) showed that multiple photosynthetic parameters demonstrated no significant differences when measured on excised leaves relative to intact leaves. Here, the only trait that did demonstrate a significant difference was stomatal conductance, which was significantly higher in excised leaves. Despite this difference, the authors showed that it was possible to use data from excised leaves to parametrise an ecophysiological model to reproduce diurnal patterns of *in situ* gas exchange. In a similar study, Akalusi *et al*., (2021) showed there to be negligible differences in light saturated rates of gas exchange and modelled estimates of photosynthetic capacity between excised and intact leaves of balsam fir (*Abies balsamea*). Moreover, the authors also demonstrated that the non-significant differences between these measurement types carried through across the growing season. In contrast to these two studies, Richardson and Berlyn (2002) focused on how spectral reflectance and chlorophyll fluorescence parameters were impacted by leaf excision in four temperate tree species. They observed that spectral and chlorophyll fluorescence changes were only minor at 12-hours post excision. Furthermore, they demonstrated that with appropriate handling to maintain leaf water status, representative data could be obtained from excised leaves as late as three days-post excision depending on the species. These studies have demonstrated the effectiveness of phenotyping traits related to photosynthesis on detached leaves of perennial woody species. As we highlight in the introduction, a similar approach is now commonly employed for many crop studies. The numerous differences that distinguish herbaceous annuals from trees suggest that the success of this approach may not carry over to crop species, thereby highlighting the timeliness of this present study. Our findings here largely provide long overdue experimental justification for applying this approach on three major crop species, representing the major monocots and dicot groups as well as C4 and C3 photosynthesis. The lack of large effects due to excision, as well as the occurrence of some small but significant effects will be discussed in more detail in the next sections.

### Stabilised leaf water potential and ABA accumulation underlie the utility of phenotyping photosynthesis in excised leaves

The most important crop species for staple foods are herbaceous annuals. A key anatomical difference between woody and herbaceous species is the composition of xylem tissue. Primary xylem is the only form of xylem for most herbaceous plants, including cereals (Růžička *et al*., 2015), although tomato can demonstrate a small amount of secondary xylem (Qi *et al*., 2020). While primary xylem comprises a relatively thin central zone within stems and leaves, secondary xylem grows out and differentiates from the vascular cambium (Kirst *et al*., 2004). Secondary xylem serves multiple purposes beyond facilitating water transport, including biomechanical support and protection against physical damage (Sperry, 2003; Morris *et al*., 2016; Miodek *et al*., 2021). Avoiding xylem damage is critical for maintaining hydraulic conductivity and protecting photosynthesis. Breakage of the water column through xylem cavitation can disrupt this. Our results are supportive of the notion that the transpiration stream of excised tomato, barley, and maize leaves is maintained following recutting them underwater as we did not observe negative effects on leaf water status measured as leaf water potential (Figure 1). The utility of performing photosynthesis on excised leaves from woody species may be supported by the presence of secondary xylem tissue, since these measurements are typically performed by excising woody branches attached to leaves of interest. Furthermore, vulnerability segmentation work has shown that branches are more resistant to xylem cavitation than terminal leaves (Zhu *et al*., 2016). Stem and leaf hydraulic conductance are coordinated (Pivovaroff *et al*., 2014), therefore excising leaves may remove the association of leaf and stem hydraulic conductance. It is plausible that the reduction in leaf water potential of intact barley leaves (Figure 1b) is a function of this and that by detaching the leaf from the stem and immersing it in water, this process increases water availability by removing the hydraulic resistance imposed by the stem. Alternatively, this difference may suggest that the plants attached to the measured leaves were marginally water stressed, however this is unlikely given the absolute water potential values were greater than -0.25MPa since we ensured that all plants were well-watered throughout growing and on the day before measurements. Moreover, the barley light-saturated gas exchange data did not demonstrate a signal of water stress in the intact leaves relative to the excised leaves (as discussed below). In general, the values for water potential obtained here and the minor difference between excised and intact leaves suggests a lack of water stress.

It has been suggested that roots are essential for stimulating ABA biosynthesis in leaves (Takahashi *et al*., 2018), however many studies that involve work with excised leaves have demonstrated that ABA levels can increase independently of roots (Bauer *et al*., 2013; Sussmilch *et al*., 2017; McAdam and Brodribb, 2018). It is therefore plausible that any treatment effects observed as a result of leaf excision could be imparted due to foliar ABA accumulation. Depending on the site and method of excision, the act of cutting a leaf or a tiller from a plant may elicit a wounding-type response. ABA can accumulate in response to wounding as a regulator of downstream defense mechanisms (Cui *et al*., 2013; Savatin *et al*., 2014). Additionally, the process of excision is likely to impair xylem tension influencing the flow rate of water to the leaf. This may also result in ABA accumulation in the leaf to stabilise leaf water status by reducing transpiration, however there no evidence presently exists to support this notion. Moreover, it is likely that the subsequent recutting of leaves underwater as per this study and other studies will quickly re-establish xylem tension by mitigating against cavitation. The role of ABA in regulating ion transport into guard cells to induce stomatal closure is well characterised (Schroeder *et al*., 2001), consequently if leaf excision were to induce leaf ABA accumulation through any basis, we could expect to see associated reductions in gas exchange as result of stomatal closure. To this end, our results demonstrated that in the morning following leaf excision there was no significant difference in the leaf ABA concentration between intact and excised leaves in either of three species (Figure 2). Performing leaf excision on the day prior to phenotyping was premeditated as we hypothesised that this would allow sufficient time for degradation of any additionally accumulated ABA. So, while our results do not allow us to determine whether excision stimulates an immediate ABA response, they are supportive of a lack of differential accumulation of ABA in leaves that are excised on the day prior to measurements. Consequently, we recommend adopting this approach when phenotyping excised leaves to avoid any confounding effects that may potentially arise due to ABA accumulation.

### Differences between excised and intact leaves for gas exchange traits are limited, but important and species-specific

Our methodology of leaf excision mirrors a widespread approach utilised by many research groups for performing photosynthesis-related phenotyping of material that is difficult to measure *in-situ* for a multitude of reasons. This approach is ultimately necessary and defines the ability to perform large scale screening of diversity that is important to characterise. Despite this, leaf excision may represent a significant shock to the leaf and plausibly elicit substantial wounding responses. Prior to this study, little data existed to support the efficiency of this approach for generating data matching that obtained from intact leaves. To this end, perhaps the most critical finding of our study is that we did not observe a significant effect of leaf excision on *A_N_*. This was true for all three species tested and across the morning and afternoon measurements (Figure 3a, e, i). This is an important finding and goes a long way to highlight the utility of this approach for assessing light-saturated photosynthesis at ambient CO_2_. While we did not observe significant treatment effects on *A_N_*, we did observe significant time of measurement effects for barley and maize, where *A_N_* declined into the afternoon (Figure 1e, i). This is indicative of the diurnal pattern of photosynthesis where, in many species, there is a decline in *A_N_* into the afternoon (Singsaas *et al*., 2000; Mohotti and Lawlor, 2002; Ainsworth *et al*., 2007; Koester *et al*., 2016; Vialet-Chabrand *et al*., 2017). This can be attributed to multiple factors including declining electron transport and changes to the apparent sink strength due to increasing leaf carbohydrate content (Singsaas *et al*., 2000; Koester *et al*., 2016). So, while this excision approach is supportive of phenotyping *A_N_*, our findings suggest that interpreting and comparing data obtained across an extended diurnal period necessitates caution regardless of whether it was collected from intact or excised leaves.

A further, and curious, significant time of day effect was detected for *c_i_* in tomato (Figure 3c). Here, *c_i_* increased into the afternoon for both treatments despite no significant changes in *A_N_* or *g_s_* that could be presupposed to lead to increases in *c_i_*. This result may be a function of lower Rubisco activity, which is suggested by declining *V_cmax_* in tomato for both treatments into the afternoon (Figure 4c). Here, lower carboxylation by Rubisco may lead to an increase in CO_2_ in the intracellular air space of the leaf in the afternoon compared to the morning, especially since *g_s_* was not observed to change across these time periods. This result highlights the importance of quantifying photosynthetic capacity to generate a broader picture of the photo-physiological state of the leaf, as opposed to relying on light-saturated measurements made under ambient CO_2_. Declining *V_cmax_* in the afternoon for tomato here may be a function of the diurnal expression patterns of Rubisco activase (Rca). Rca is the molecular chaperone that defines Rubisco activity, and its expression has been demonstrated to decline significantly into the afternoon (Perdomo *et al*., 2021). The afternoon decline in tomato *iWUE* (Figure 3d) mirrors the increase in *c_i_* (Leakey *et al*., 2019) and is a function of the combination of the decline in *A_N_* (Figure 3b) and increase in *g_s_* (Figure 3c), despite those changes being non-significant in isolation.

One of the more surprising results from our study was that maize was the only one of the three species where we detected a treatment effect on *g_s_* (Figure 3j). Here, the excision process resulted in a significant decline in *g_s_* compared to intact leaves. This result is somewhat surprising since maize leaves are larger and appear more physically robust than barley leaves and the petiole of tomato leaves. Nevertheless, we ascribe confidence to this result since it was consistent across the two time points, therein highlighting the impairing influence of leaf excision on stomatal opening in maize. The C_4_ carbon concentrating mechanism that defines the relative de-coupling of *g_s_* and *A_N_* in C_4_ species relative to C_3_ species (Leakey *et al*., 2019) may underly why we do not observe a concurrent treatment effect on *A_N_* in maize. Namely, the decrease in *g_s_* observed in the excised leaves was not sufficient to decrease *c_i_* enough to lose CO_2_ saturation of *A_N_* (Figure k) as evident from the *A_N_*/*c_i_* responses (Figure 3j). The treatment effect of leaf excision on *g_s_* in maize but not on *A_N_*, led to expected results for *c_i_*, where it decreased significantly at both time points due to decreased capacity for CO_2_ uptake (Figure 3k). Concurrently, the stomatal resistance to water loss was greater, such that *iWUE* was higher at both time points and more so again in the afternoon (Figure 3l). Taken together, these results show that the time-of-day effects on light-saturated rate of gas exchange are stronger in intact maize leaves, thereby highlighting the potential for longer measurement days when working with excised leaves.

Despite the above-described differences observed in maize, we did not observe any significant effects in terms of treatment or time on the photosynthetic capacity traits, namely *V_pmax_* and *V_max_* (Figure 2k, l). This suggests that reduced *g_s_* at the starting point of the *A_N_*-*c_i_* response measurements, i.e., ambient CO_2_, in excised leaves did not compromise the ability to assess the rate of photosynthetic induction from low-to-ambient CO_2_ concentrations or the rates of photosynthesis when *g_s_* is minimal at high CO_2_ concentrations. Our results are therefore supportive of performing *A_N_*-*c_i_* response curves in excised maize leaves to model photosynthetic capacity. In general, our results are also supportive of performing *A_N_*-*c_i_* response curves on excised leaves of the C_3_ species. We did detect significant reductions in *V_cmax_* (barley; Figure 4g) but the difference was minor and not exaggerated by the interaction with time.

### Reflectance and chlorophyll fluorescence appear relatively unchanged between intact and excised leaves

There were no statistically significant differences detected for leaf reflectance between the treatments for any of the species on a wavelength-by-wavelength basis (Figure 5). Despite this, the pattern of p-values across the spectrum were surprisingly unique between the species and different between the time points for barley. In general, if we consider the dominant factors that influence leaf reflectance (Kokaly *et al*., 2003), the p-value patterns suggest the most excision-sensitive aspects of reflectance were due to leaf water content in tomato and light scattering in the mesophyll in barley and maize (Figure 5). The results from barley and maize here are similar to those observed across certain tree species by Richardson and Berlyn, (2002), where the authors and others suggest reductions in near-infra red spectra reflectance in excised leaves are related to leaf structural changes at the cellular level. Here, declining leaf water status results in cytoplasmic shrinkage, such that light scattering alters at the interface between cell walls and water (Carter, 1991; Aldakheel and Danson, 1997; Richardson and Berlyn, 2002). However, we do not observe perturbed leaf water status in excised leaves (Figure 1), consequently this may reflect why the patterns we highlight in terms of differences in reflectance between the treatments do not pass significance thresholds (Figure 5). The study of Richardson and Berlyn, (2002) tested differences between excised and intact leaves over several days post-excision and did not begin to detect significant differences in reflectance indices and chlorophyll fluorescence until 48 hours after excision. Consequently, it is possible that we may have observed differences in reflectance and other traits had we measured for a further day, however this is not common practice with crop species, and we would advise against this.

To resolve any potential reflectance differences between intact and excised leaves, we calculated five commonly employed reflectance indices (Supplemental Table S1, Supplemental Figures S3-5). In doing so, we were only able to detect two statistically significant differences between intact and excised leaves. For barley, the MSI was significantly higher in the morning in excised leaves suggesting a reduction in leaf water content, however this effect was not apparent in the afternoon (Supplemental Figure S4d). In maize, a significant treatment effect on the Datt index, which is positively correlated to chlorophyll content (Lu *et al*., 2015), suggested that chlorophyll content was reduced in excised leaves (Supplemental Figure S5b). Here, it could be hypothesised that ABA accumulation would be an the underlying mechanism resulting in this difference, since there is also some evidence linking leaf yellowing and reductions in leaf chlorophyll to ABA, however that is evidently not the situation in our study (Figure 2) and this is typically a longer-term effect (Wang *et al*., 2018). In general, our results here suggest that reflectance changes between excised and intact leaves are slight, which is similar to what has previously been shown in woody species (Richardson and Berlyn, 2002).

Our results also mirror those of Richardson and Berlyn, (2002) in that we saw no differences in *F_v_*/*F_m_* in either of methods through which it was assessed in this study (Figure 6 and Supplemental Figure S6). This highlights how electron flow in PSII is not significantly impacted by this approach, so there is no impairment on the photosynthetic apparatus. In our study we also demonstrated no significant differences in the dynamics of NPQ, expect for a minor reduction in the rate of induction in excised tomato leaves when measured using the gas exchange leaf chamber fluorometer (Supplemental Figure S6a). The fact that this was not detected the following day using the closed chlorophyll fluorescence system suggests that this is more likely to result from insufficient time allowed for dark adaption, especially since *F_v_*/*F_m_* was also bordering on being significantly reduced in excised leaves when measured in the leaf chamber fluorometer also (p = 0.07; Supplemental Figure S6a).

## Conclusion

Screening large amounts of diversity for photosynthesis-related traits is only realistically possible with excised leaves. Therefore, the evidence presented in this study that validates this approach, in terms of its affiliation to intact-leaf measurements, is welcomed. Despite this, our study does highlight the existence of specifies-specific treatment effects, e.g., *g_s_* reduction in maize, that the community should be aware of. Moreover, this should encourage those who are planning on adopting this approach to validate if it is suitable for their species of interest. One salient point that our study does not address is the potential for differential responses to excision on an intra-specific basis. An obvious further extension to this study would be to test how genetically distinct genotypes within a species perform following leaf excision as this has the potential to have large consequences for studies of this nature.

## Supplemental Material

Supplemental Figure S1. Overview of experimental workflow.

Supplemental Figure S2. Mean hyperspectral reflectance data. The solid line represents the mean reflectance, and the shaded area represents the standard error of the mean. (a) Tomato AM (b) Tomato PM (c) Barley AM (d) Barley PM (e) Maize AM (f) Maize PM

Supplemental Figure S3. Hyperspectral reflectance derived indices for tomato. (a) Red Edge- Normalised Difference Vegetation Index (RE-NDVI), (b) Datt Index (c) Normalised Difference Nitrogen Index (NDNI), (d) Moisture Stress Index (MSI), and (e) Leaf Water Index (LWI)

Supplemental Figure S4. Hyperspectral reflectance derived indices for barley. (a) Red Edge- Normalised Difference Vegetation Index (RE-NDVI), (b) DATT Index (c) Normalised Difference Nitrogen Index (NDNI), (d) Moisture Stress Index (MSI), and (e) Leaf Water Index (LWI)

Supplemental Figure S5. Hyperspectral reflectance derived indices for maize. (a) Red Edge- Normalised Difference Vegetation Index (RE-NDVI), (b) DATT Index (c) Normalised Difference Nitrogen Index (NDNI), (d) Moisture Stress Index (MSI), and (e) Leaf Water Index (LWI)

Supplemental Figure S6. Boxplots showing variation in chlorophyll fluorescence associated traits between excised and intact leaves measured using the leaf chamber fluorometer.

Supplemental Table S1. Reflectance indices used in this study.

## Acknowledgments

This work was supported by the European Union’s Horizon2020 research and innovation programme (No.862201) project CAPITALISE. We thank Dr Cristina Sales and Dr Angie Burnett for assistance in generating the ABA data. We thank Dr Sara Lopez-Gomollon for providing the tomato seed. We thank Katie Shaw for assistance in performing the modelling of the chlorophyll fluorescence data and Prof Howard Griffiths and Dr Jessica Royles for assistance with the leaf water potential measurements. For the purpose of open access, the authors have applied a Creative Commons Attribution (CC BY) licence to any Author Accepted Manuscript version arising from this submission.

## Author contributions

JNF, TL and JK designed the study. JNF and TJ carried out the experiments. JNF and TJ performed the data analyses. All authors helped to interpret the results. JNF and JK wrote the paper with input from all the authors.

## Abbreviations

ABA: Abscisic acid
*A_N_*: Net photosynthetic CO_2_ assimilation
*c_i_*: Intracellular CO_2_ concentration
*F_m_*: Dark adapted maximum fluorescence
*F_m_’*: Maximum fluorescence
*F_v_*/*F_m_*: Maximum photosystem II efficiency
*g_s_*: Stomatal conductance
*iWUE*: Intrinsic water use efficiency
*J_max_*: Maximum rate of electron transport for RuBP regeneration
LWI: Leaf Water Index
MSI: Moisture Stress Index
NDNI: Normalised Difference Nitrogen Index
NPQ: Non-photochemical quenching
PAR: Photosynthetically active radiation
RE-NDVI: Red Edge-Normalised Difference Vegetation Index
*V_cmax_*: Maximum rate of Rubisco carboxylation
*V_pmax_*: Maximum rate of PEPc carboxylation
*V_max_*: Maximum rate of photosynthesis

