## Supplementary figures and images for "Leaf excision has minimal impact on photosynthetic parameters across crop functional types"

### supplementary figure 1

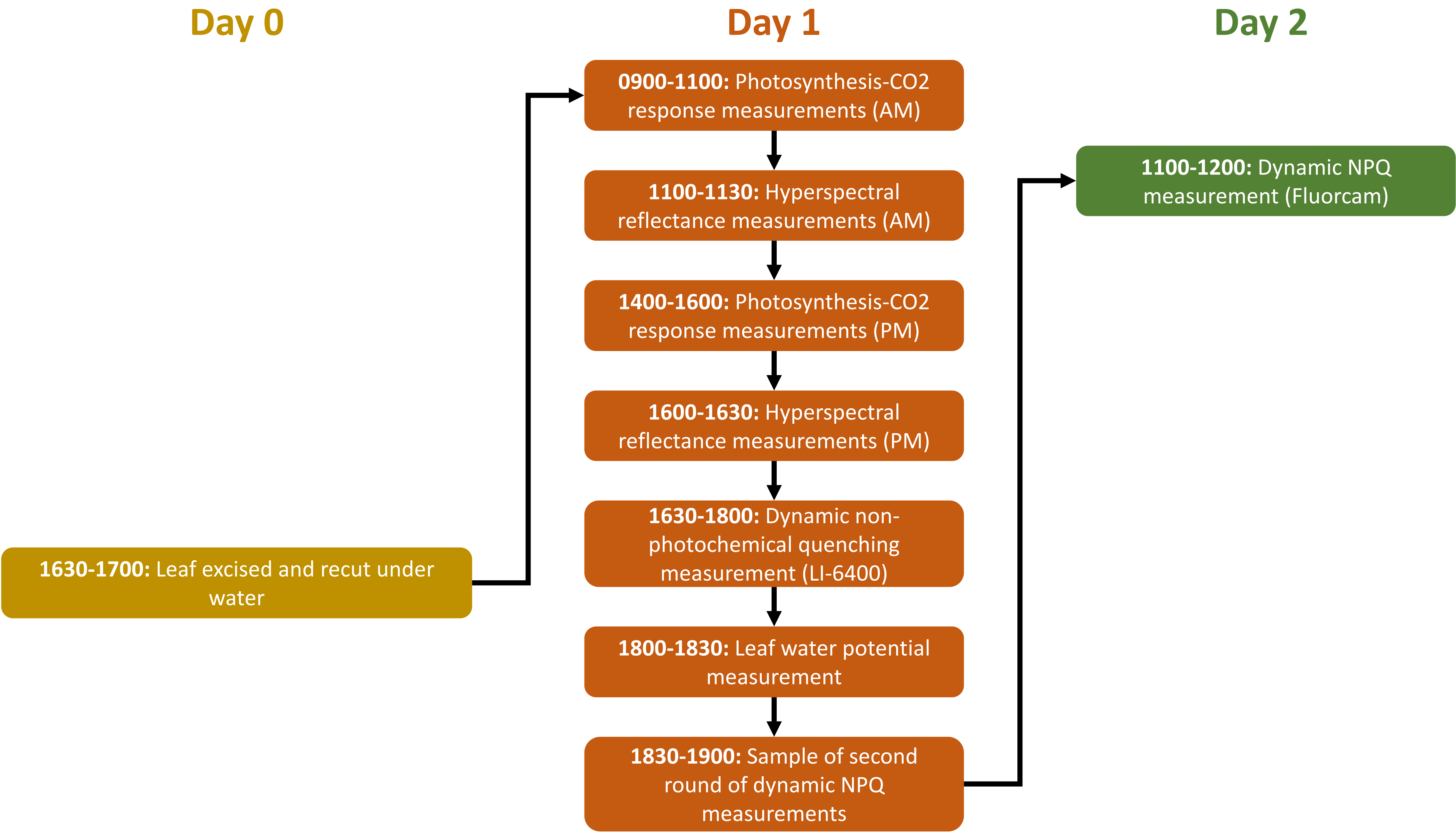

### supplementary figure 2

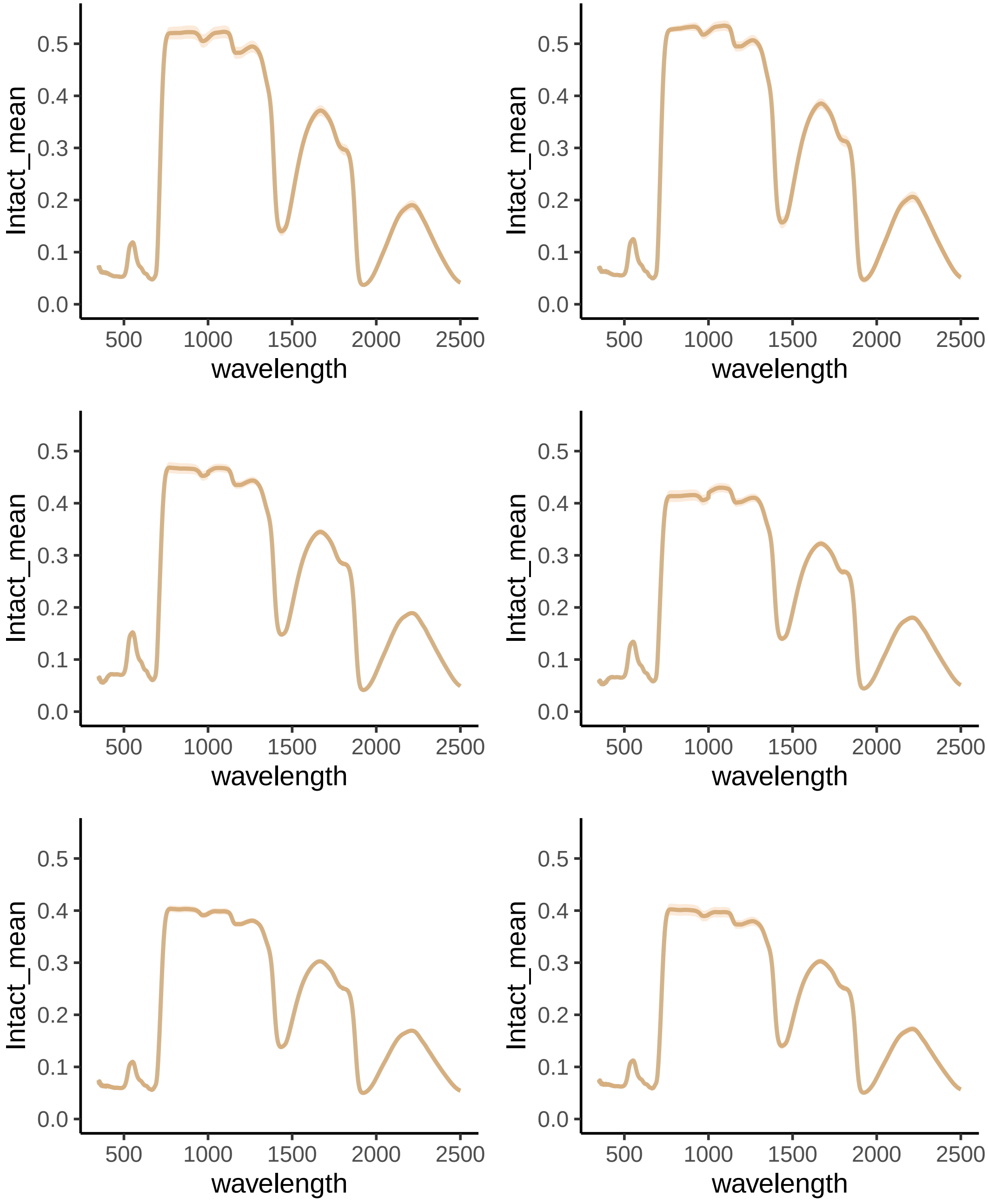

### supplementary figure 3

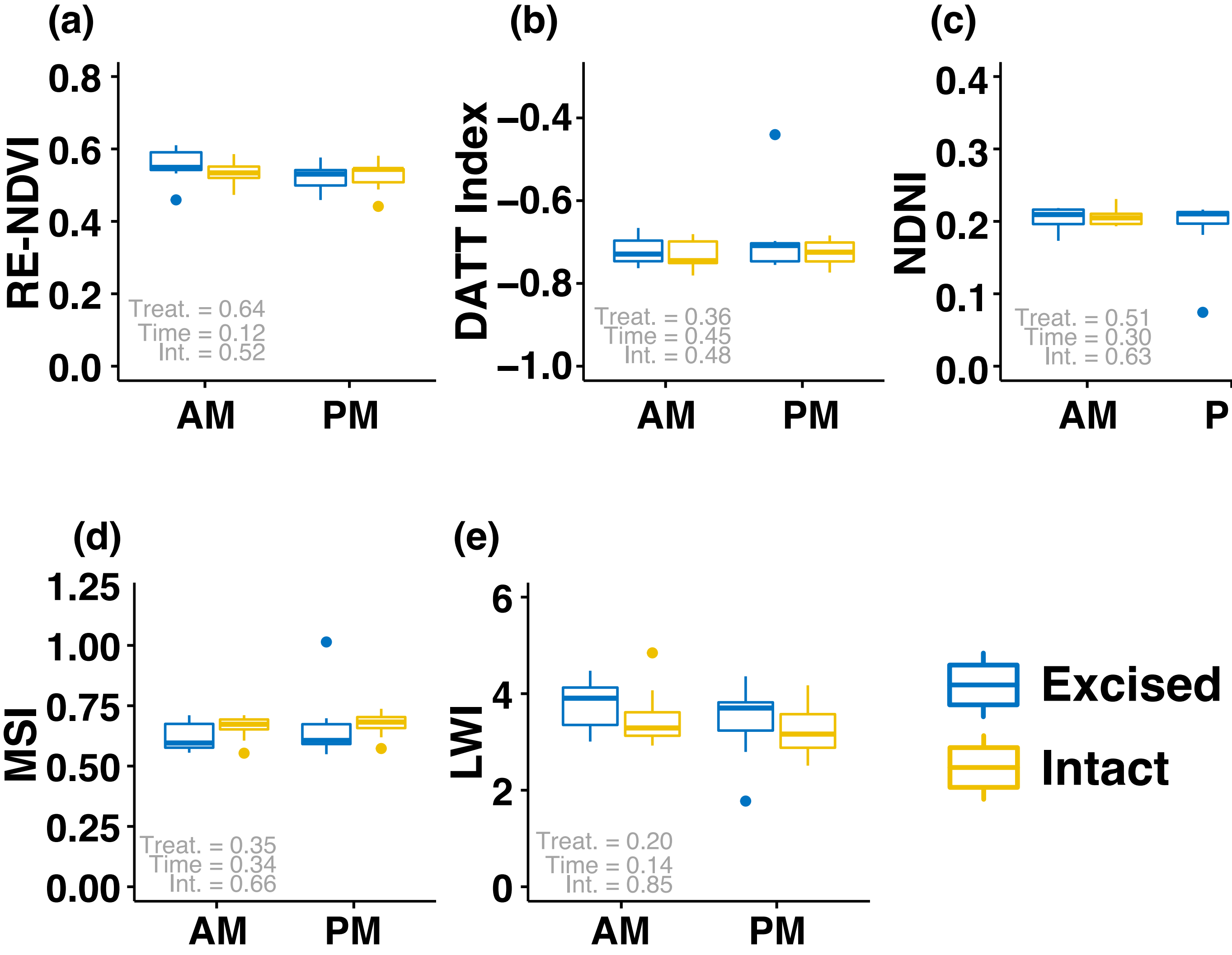

### supplementary figure 4

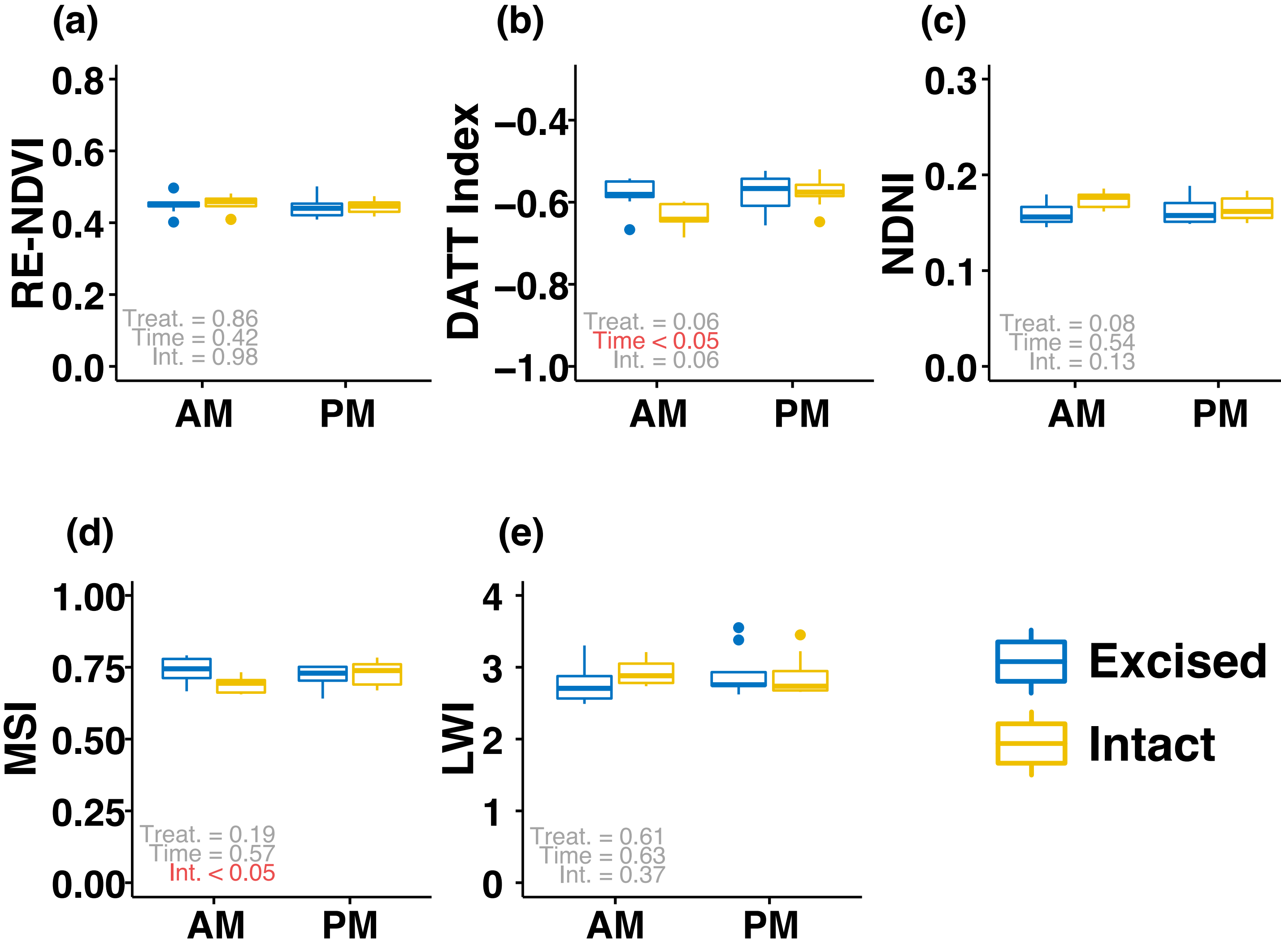

### supplementary figure 5

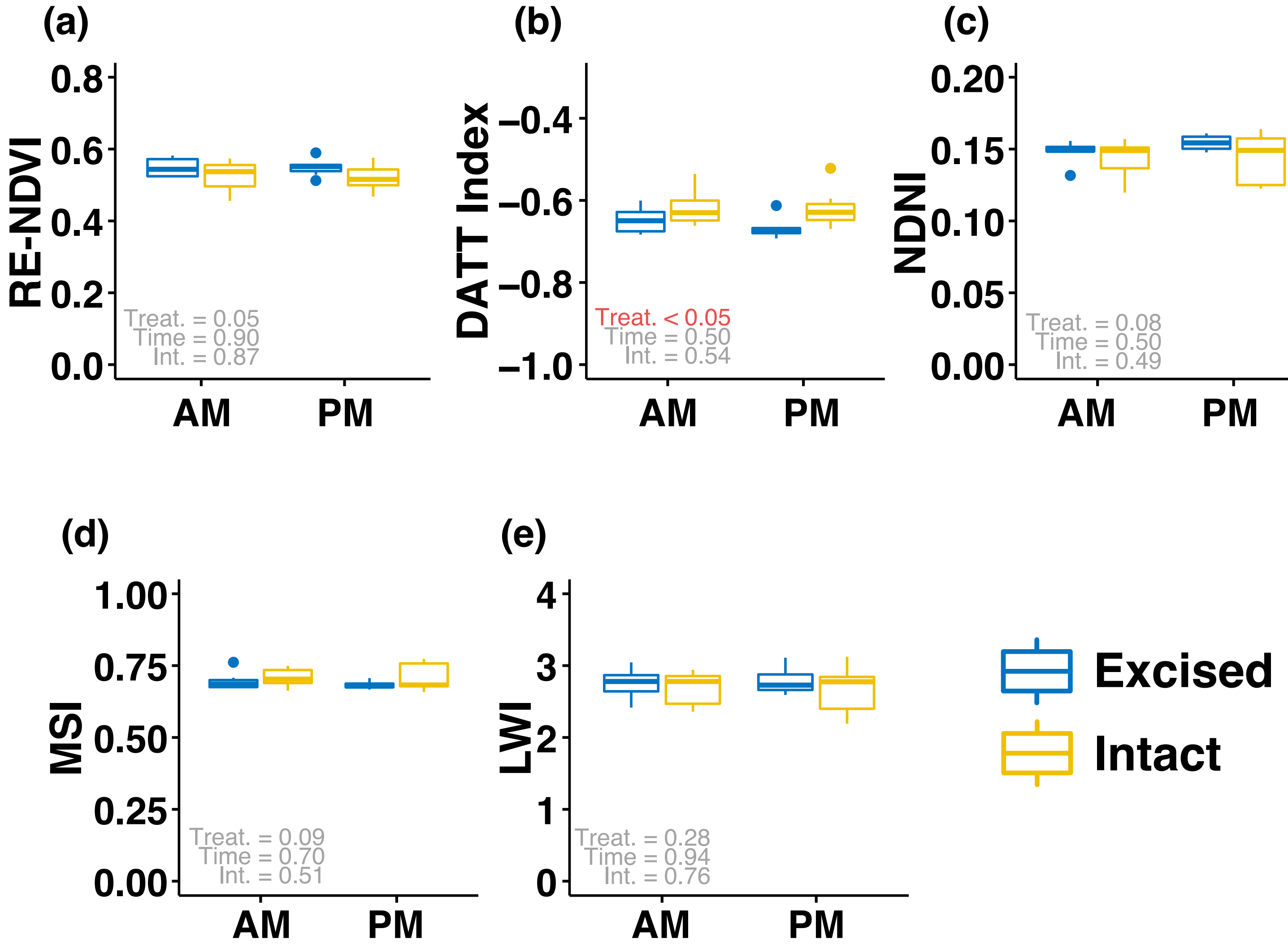

### supplementary figure 6

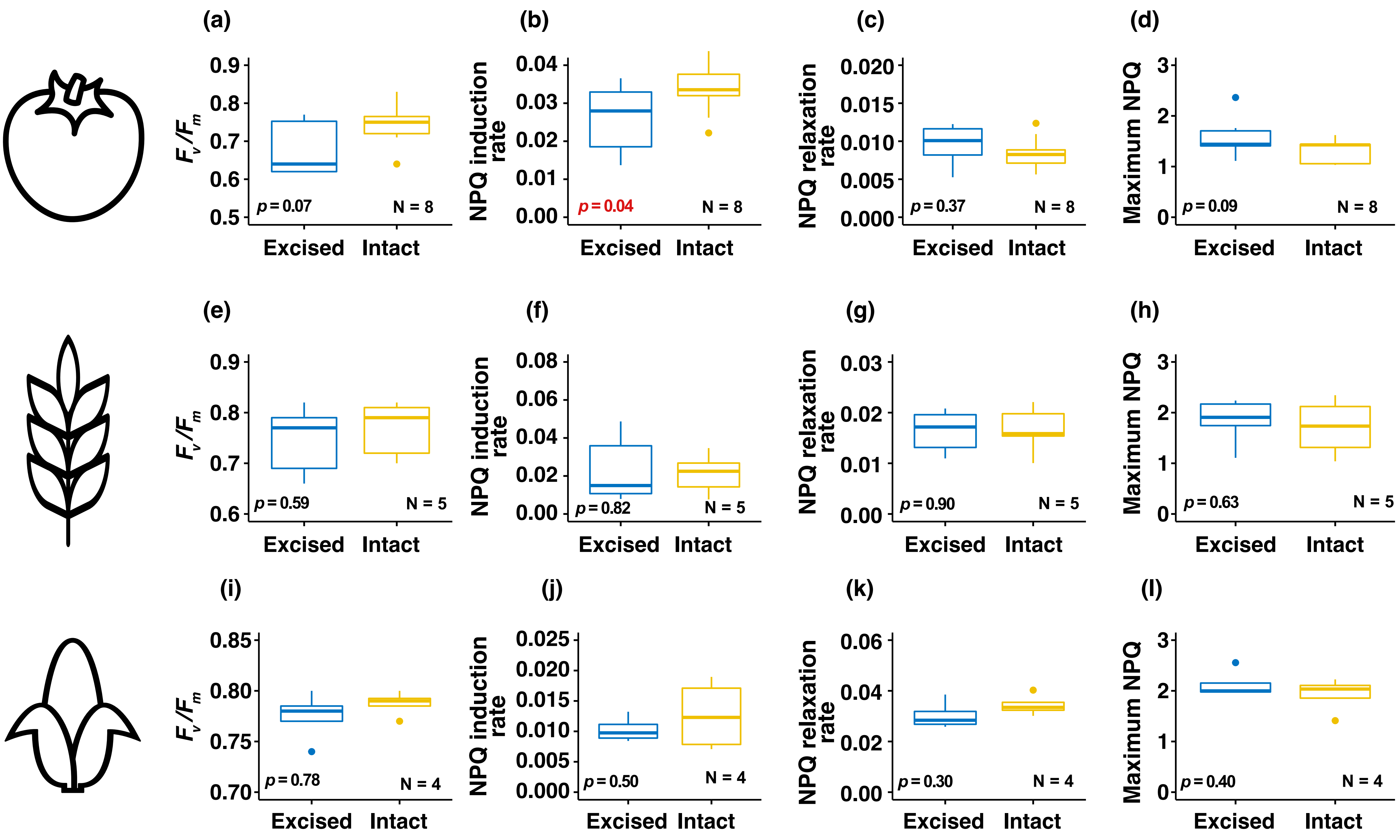
